# N^6^-methyladenosine primes the malaria parasite for transmission

**DOI:** 10.1101/2025.03.28.645929

**Authors:** Selina Mussgnug, Tiziano Vignolini, Justine E. Couble, Patty Chen, Dayana Farhat, Ameya Sinha, Grégory Doré, Delia Dupré, Marion Lambault, Louisa Hadj Abed, Peter R. Preiser, Peter C. Dedon, Jessica M. Bryant, Sebastian Baumgarten

## Abstract

Sudden environmental changes are a recurring challenge for unicellular organisms, but a necessity for many to progress through their lifecycle. To transmit from its human host to mosquito vector, malaria parasites differentiate into male and female, semi-quiescent stages that can re-initiate development within seconds after transmission. Here, we identify the RNA modification N^6^-methyladenosine (m^6^A) as the mediator of a rapid, sex-specific, and temperature-sensitive mechanism to restructure protein synthesis during transmission. We find that male parasites maintain high levels of translation during their semi-quiescence that are rapidly repressed following mosquito uptake. This translational shutdown is essential for the continuation of male parasite development and depends on the m^6^A-binding protein YTH.2. We further show that m^6^A and YTH.2 are already present prior to transmission, but that their repressive interaction requires a temperature drop accompanying the exit from the human host. Hence, m^6^A appears to prime the parasite transcriptome and subsequently converts an environmental shift into a rapid translational response.

## INTRODUCTION

All living organisms evolved to thrive within the physical boundaries of their ecosystem, yet sudden environmental changes are a recurring challenge for many, especially unicellular organisms. In some cases, such fluctuations can coincide with a switch in the developmental program of an organism, for example the entry into a dormant stage whose purpose is to withstand unfavorable conditions^1–6^. On the other hand, many eukaryotic parasites rely on the transmission to an intermediate vector to be able to infect new hosts. A particularly extreme example is the human malaria parasite, *Plasmodium falciparum*. During the infection within the human host, the parasite replicates within red blood cells (RBC) over the course of a ∼48h asexual replicative cycle, before up to 32 daughter cells erupt from the RBC and infect new ones^7^. To be taken up by an Anopheles mosquito vector, however, malaria parasites can exit this asexual replicative cycle^8–10^ and develop over the course of ∼10 days into male and female gametocytes, the human-to-mosquito transmission stages (Figure 1A). Male and female gametocytes do not replicate and enter a semi-quiescent state that is characterized by overall low metabolic activity. Although gametocytes cannot re-enter the asexual replicative cycle, they can remain quiescent yet transmissible for several days^11^.

**Figure 1:**
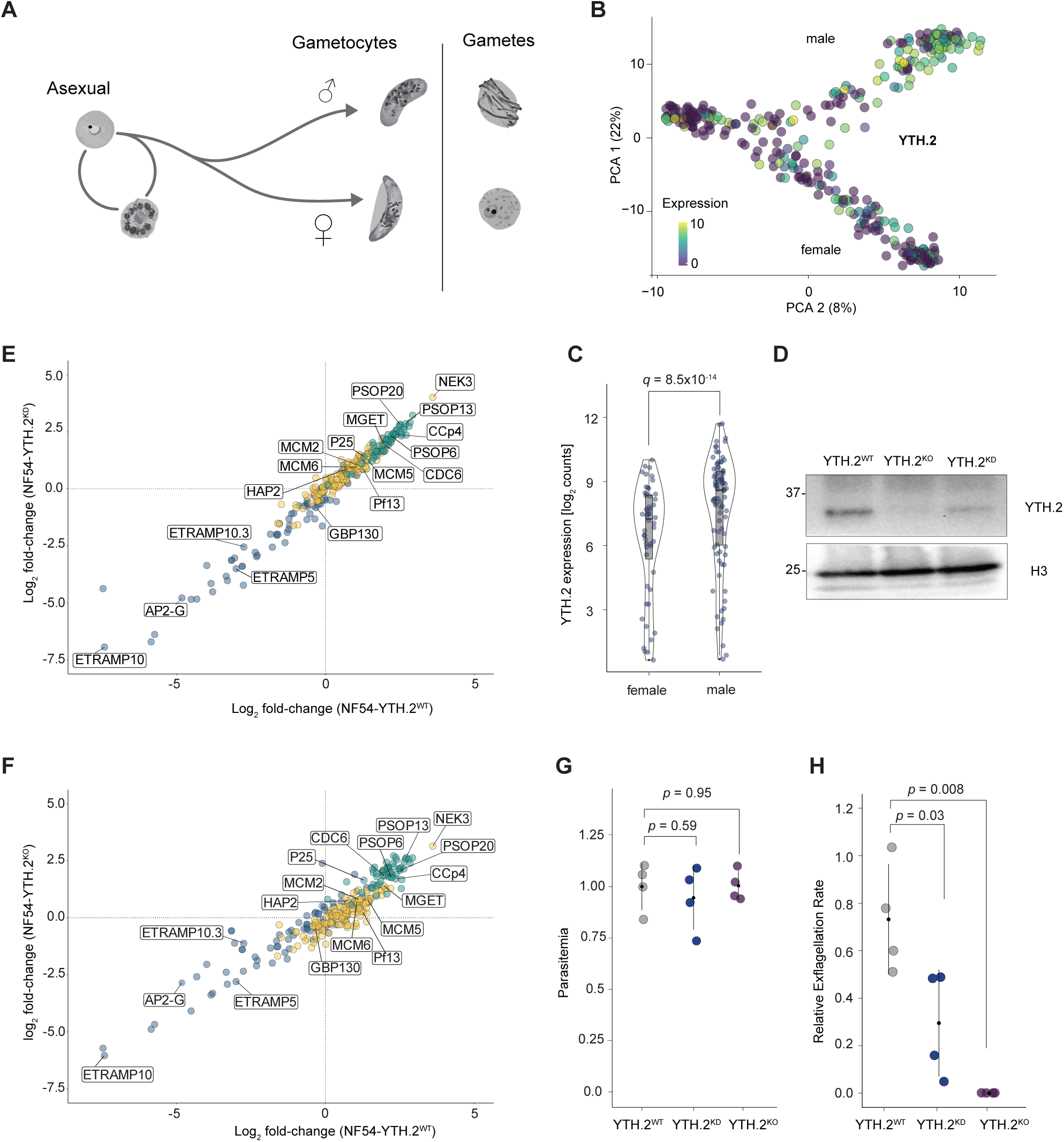
The m^6^A reader YTH.2 is essential for male parasite transmission. (A) Life cycle diagram of the developmental transition from asexually replicating *P. falciparum* to the formation and bifurcation of semi-quiescent, male and female gametocytes. Following transmission (indicated by the black line), development is re-initiated to form micro- and macrogametes. (B) Expression levels of YTH.2 in individual cells as determined by Smart-Seq2^59^ during sex determination and differentiation of *P. falciparum* gametocytes. (C) Comparison of YTH.2 expression levels^59^ of YTH.2 in male and female gametocytes. (D) Western Blot of YTH.2 protein in mature gametocytes of NF54-YTH.2^WT^ (left), NF54-YTH.2^KO^ (center) and NF54-YTH.2^KD^ (right) parasites. Proteins were collected ten days post gametocyte induction. Molecular weights (in kDa) are indicated on the left. Histone H3 serves as loading control. (E) Comparison of marker gene expression changes of immature NF54-YTH.2^WT^ parasites (sampled at day 4 post gametocyte induction) to NF54-YTH.2^WT^ (abscissa) and NF54-YTH.2^KD^ (ordinate). Immature (blue), male (yellow) and female (green) marker genes were taken from ref.^42^ (F) Same as for (E), but for NF54-YTH.2^KO^ (G) Relative numbers of viable NF54-YTH.2^WT^, NF54-YTH.2^KD^ and NF54-YTH.2^KO^ mature gametocytes. *p-*values are calculated with a Welch-two sample t-test. Black dot: mean; vertical line: standard deviation of the mean (SDM). (H) Exflagellation rates of NF54-YTH.2^WT^, NF54-YTH.2^KD^ and NF54-YTH.2^KO^ parasites. *p-*values are calculated with a Welch-two Sample t-test. Black dot: mean; vertical line: SDM.

The rapid re-initiation of development from gametocytes into gametes (i.e. ‘activation’) is triggered by, and relies on, the sensing of multiple individual environmental changes that occur during host-to-vector transmission^12^. The activation of the semi-quiescent gametocyte state to develop into gametes requires an increase in pH that occurs in the mosquito midgut during the blood feed and xanthurenic acid (XA), a product of tryptophan oxidative metabolism, which amplifies the effect of the pH increase^13^. In addition, a further drop in environmental temperature is necessary to complete differentiation into gametes, but not for activation of gametocytes itself^14^.

However, the immediate developmental trajectory following the uptake by a mosquito vector substantially differs between the two gametocyte sexes. Female gametocytes egress from the red blood cell and differentiate into a macrogamete that is characterized by the export of a range of proteins to its cell surface^15,16^. In contrast, male gametocytes undergo three rounds of rapid genome replication and develop into a total of eight elongated, motile microgametes within less than ten minutes. This process of ‘exflagellation’ is triggered within seconds following mosquito uptake and signaled through a multitude of secondary messengers^14^ and kinases that facilitate the developmental switch^17,18^.

For female gametocytes, one mechanism that is key to the preparation for successful transmission is the transcription, yet extensive translational repression of mRNAs whose products are only required once the parasite is taken up by the mosquito vector^19–21^. This translational repression is mediated by an RNA-binding complex centered around the DDX6 RNA helicase (‘DOZI’)^20,22–27^. It has been hypothesized that this repression is maintained by the formation of P-body-like compartments within female cells that keep transcripts inaccessible to ribosomes^21,25^. In addition, changes in secondary mRNA structure have been found to (de-) repress protein synthesis following transmission^28^.

Conversely, male gametocytes do not seem to engage in extensive transcript stockpiling/translational repression in preparation for host-to-vector transmission^19^. Thus, it is unclear if and how they can prepare for a much faster developmental progression immediately after transmission compared to female gametocytes. In addition, despite evidence supporting post-transcriptional regulation as central for the female parasite’s preparation for transmission, the mechanisms by which any mRNA is specifically recognized for translational control or the dynamics of protein synthesis upon mosquito uptake of either sex on a transcriptome-wide level remain currently unclear.

One mechanism that can specifically target an mRNA transcript for distinct post-transcriptional regulation is the methylation of adenosine at the sixth nitrogen position (i.e. N^6^-methyladenosine, m^6^A)^29–32^. *P. falciparum* features some of the highest m^6^A methylation levels recently measured^33^, and its genome encodes two specific ‘reader’ proteins that contain an m^6^A-binding YT521-B domain (YTH.1 and YTH.2)^33–35^. YTH.1 features a domain architecture conserved amongst apicomplexans and plants and functions in the 3’-end processing of m^6^A-methylated transcripts^36–38^. In contrast, YTH.2 has no clear homology to any m^6^A-binding proteins outside *Plasmodium sp.* and is essential for the asexual replicative cycle of *P. falciparum,* possibly by modulating translation of cognate m^6^A-methylated transcripts^39^. However, among all developmental stages in the human host, YTH.2 is most highly expressed in gametocytes, suggesting a vital role in parasite transmission^33^.

Despite the evidence that precise control of protein synthesis is essential for efficient parasite transmission, the transcriptome-wide dynamics of translation immediately before and after the re-initiation of male and female parasite development are still unclear. We therefore set out to investigate how the parasite can anticipate the sudden exit from its semi-quiescent gametocyte stage and how m^6^A mRNA methylation can help reshape the parasite transcriptome during the developmental transition that occurs during host-to-vector transmission. We find that YTH.2 deletion does not affect the development into mature gametocytes, but specifically inhibits the developmental progression of male gametocytes into gametes after mosquito uptake. Combining quantitative, nucleotide-resolution m^6^A sequencing with single-cell, RNA-protein proximity labelling, we find that YTH.2 is predominantly expressed in male gametocytes and binds to m^6^A-methylated transcripts. We show that male parasites feature an opposite translational program compared to females: extensive protein synthesis in mature gametocytes followed by rapid translation repression after the re-initiation of development into gametes. We further find that this translational shutdown of m^6^A-methylated transcripts depends on YTH.2. In addition, while m^6^A and YTH.2 are already abundantly present in male gametocytes before transmission, we show that their interaction increases ten-fold upon temperature decrease, suggesting that the environmental change accompanying parasite transmission facilitates their repressive function. Hence, m^6^A appears to prime the male gametocyte transcriptome for a rapid translational rewiring and facilitates a post-transcriptional, environment-dependent ‘paternal-to-zygote’ transition.

## RESULTS

### The m^6^A reader protein YTH.2 is essential for male gametocyte transmission

m^6^A mRNA methylation has become a strong candidate for a potential regulatory mechanism during *P. falciparum* transmission because during development in the human host, its reader protein YTH.2 reaches peak expression during gametocyte maturation^33^. Resolving its expression pattern in a sex-dependent manner using previously published single-cell RNA sequencing (scRNA-seq) data shows that YTH.2 is predominantly expressed in male gametocytes (Figure 1B). Of all cells sampled, YTH.2 could be detected in 76% (n = 84/110) of male cells, but in only 41% (n = 52/126) of female cells. In addition, expression levels in cells that do express YTH.2 are significantly lower in female than in male cells (Figure 1C), suggesting a possible sex-specific role of YTH.2 during *P. falciparum* host-to-vector transmission. In contrast, YTH.1 is detected equally in male and female gametocytes and at similar expression levels.

YTH.2 is essential for the asexual replicative cycle^39,40^; therefore, we used an inducible knockout strategy to functionally characterize YTH.2 in transmission stages. Using CRISPR/Cas9, we introduced *loxP* sites up- and downstream of the endogenous genomic *yth.2* locus in a strain background that expresses a dimerizable Cre recombinase^41^. We then induced the deletion of the entire *yth.2* locus either at the onset of gametocytogenesis (0 days after gametocyte induction) or during early gametocytogenesis (4 days after gametocyte induction). Interestingly, deletion of *yth.2* at either time point did not phenotypically affect gametocyte maturation, with mature stage V gametocytes being readily detectable. Western blot analysis of stage V gametocytes at day 10 post gametocyte induction showed that YTH.2 protein was readily detectable in untreated cells (NF54-YTH.2^WT^), but was undetectable when the gene was deleted at Day 0 (NF54-YTH.2^KO^) and substantially reduced when deleted at Day 4 of gametocytogenesis (NF54-YTH.2^KD^, Figure 1D).

To further confirm that YTH.2 does not affect gametocyte maturation of either sex, we compared mRNA levels between immature NF54-YTH.2^WT^ gametocytes and mature NF54-YTH.2^WT^, NF54-YTH.2^KD^, and NF54-YTH.2^KO^ gametocytes. For either NF54-YTH.2^KD^ (Figure 1E) or NF54-YTH.2^KO^ (Figure 1F), the signature of up- and down-regulated marker genes of early and late transmission stage development^42^ is identical to that of NF54-YTH.2^WT^ parasites, further indicating that YTH.2 deletion does not disrupt the transcriptional program underlying gametocytogenesis. In addition, direct quantification of viable, mature parasites at day 10 of gametocytogenesis did not reveal any significant difference between parasites with an intact *yth.2* locus and those with *yth.2* deletion at day 0 or at day 4 (Figure 1G). We also did not find a change in the ratio of male and female cells. These data indicate that YTH.2 is not essential for gametocyte maturation, and that YTH.2 protein expressed early in gametocytogenesis is stable and remains present throughout transmission stage development.

Next, we investigated the effects of YTH.2 perturbation on the re-initiation of development following transmission *in vitro*. While YTH.2 perturbation did not affect differentiation of mature female gametocytes into macrogametes, NF54-YTH.2^KD^ showed a significant decrease in mature male gametocyte differentiation into microgametes. Moreover, no male gametes were observed for NF54-YTH.2^KO^ parasites (Figure 1H). These data indicate that while YTH.2 does not affect the maturation of *P. falciparum* into mature male and female gametocytes on a phenotypic or transcriptomic level, YTH.2 may play a specific role one developmental step later as male gametocytes reinitiate development into microgametes.

### Orthogonal mapping of m^6^A during *P. falciparum* gametocyte development

In contrast to YTH.2, MT-A70, the central protein of the *P. falciparum* m^6^A methyltransferase complex, is expressed in both male (78/110 cells) and female (73/126) gametocytes at similar levels (Figure 2A,B). Moreover, m^6^A itself was readily detected on gametocyte mRNA using LC-MS/MS at levels similar to those measured in asexually replicating cells^33^ (Figure 2C). Thus, we wanted to determine any gametocyte sex-specific patterns with regard to m^6^A and mapped m^6^A sites transcriptome-wide at day 7 and 10 post gametocyte induction. To resolve m^6^A modification sites at single-nucleotide resolution with the limited input amounts that can be retrieved from gametocytes, we developed an antibody-based approach termed m^6^A-eCLIP2, which combines cross-linking m^6^A pulldowns^43–45^ with multiplexed input samples^46^. Here, multiple samples are directly barcoded using specific RNA adapters, allowing for a single pool to be processed, which increased overall RNA amounts and eliminated technical biases. m^6^A methylation sites are then identified based on cross-linking an m^6^A antibody to the methylated adenosine. During library preparation, the covalent antibody-m^6^A interaction can lead to a nucleotide substitution (most often cytosine to uridine, C-to-U) by the reverse transcriptase at the first position downstream of the m^6^A site (Figure 2D)^44^.

**Figure 2:**
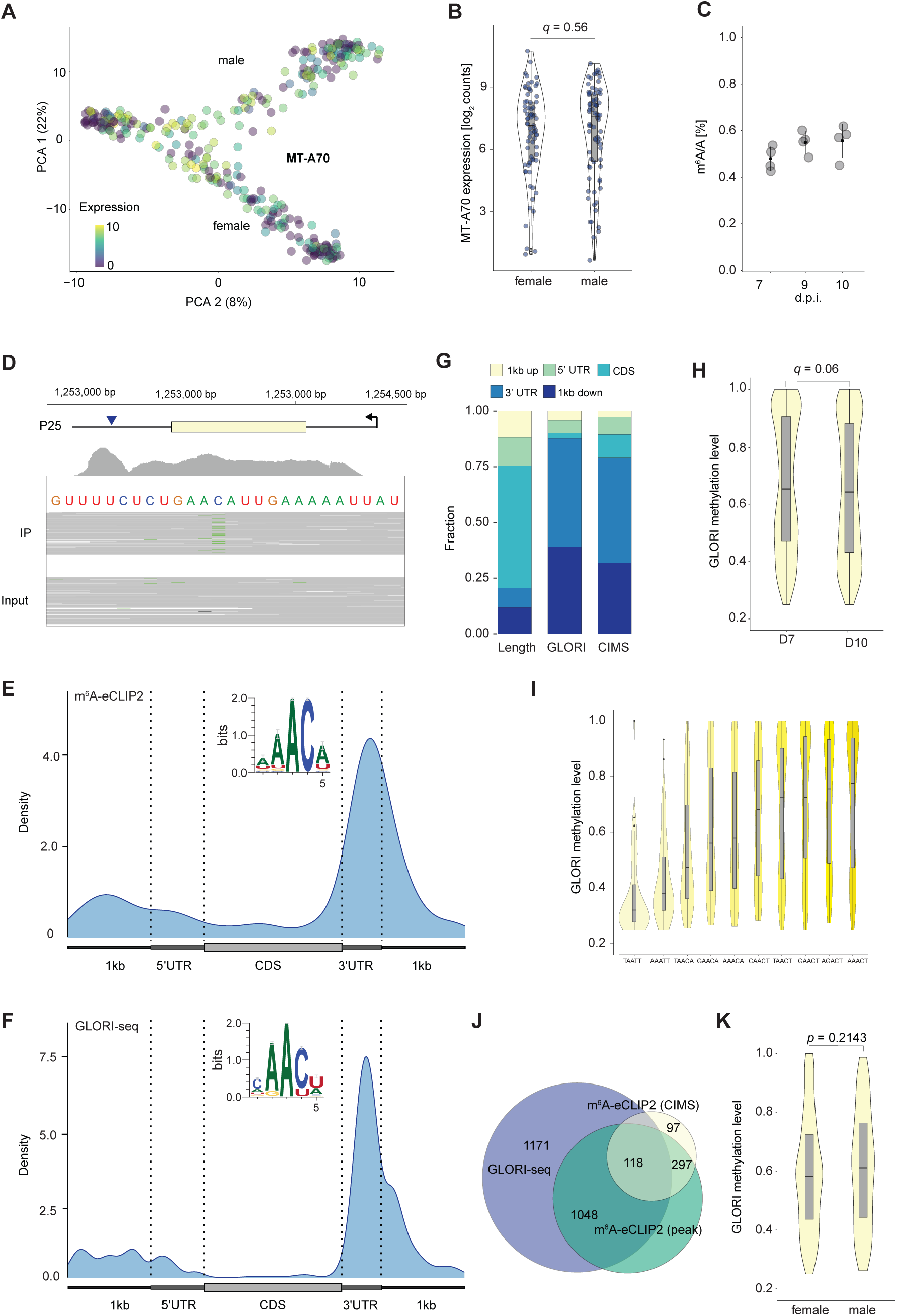
Orthogonal mapping of m^6^A during *P. falciparum* gametocyte development. (A) Expression levels of MT-A70 in individual cells as determined by Smart-Seq2^59^ during sex determination and differentiation of *P. falciparum* gametocytes. (B) Comparison of gene expression levels^59^ of MT-A70 in male and female gametocytes. (C) Percentage of N^6^-methyladenosine (m^6^A) of all canonical adenosines (A) at three timepoints of gametocyte development. d.p.i.: Days post gametocyte induction. (D) Example of a m^6^A site detected by m^6^A-eCLIP2 in the mRNA transcript encoding P25. C-to-U conversion at the -1 position of the putatively modified adenosine are indicated (green) within individual aligned reads (grey). and only detectable in the crosslinked immunoprecipitated sample (IP, top), but not the un-crosslinked control sample (Input, bottom). Note the increased read coverage of the crosslinked, immunoprecipitated sample at the site of the CIMS. (E) Metagene plot representing the global distribution of m^6^A CIMS within transcripts detected by m^6^A-eCLIP. Insert: Nucleotide motif surrounding the m^6^A site and the CIMS. (F) Metagene plot representing the global distribution of m^6^A sites within transcripts detected by GLORI-seq. Insert: Nucleotide motif surrounding the m^6^A site. (G) Fractions of different mRNA transcript regions according to total length within the mRNA transcriptome (left), and the fraction of m^6^A sites identified in each region either by GLORI-seq (middle) or m^6^A-eCLIP2 (right). (H) Comparison of m^6^A methylation levels as determined by GLORI-seq at the two different time points of gametocytogenesis. *p-*value calculated using an unpaired Wilcoxon Rank Sum test (I) m^6^A methylation levels as determined by GLORI-seq classified by the surrounding nucleotide motif (J) Overlap of high-confidence m^6^A sites identified by m^6^A-eCLIP2 (CIMS + peak calling) and GLORI-seq (see Materials and Methods). (K) Comparison of m^6^A methylation levels as determined by GLORI-seq between female- and male-dominant transcripts^42^. *p-*value was calculated using unpaired Wilcoxon Rank Sum test.

Using this approach, we identified a total of 244 and 241 crosslink-induced mutation sites (CIMS) from gametocytes collected at day 7 and day 10, respectively, that feature a significant increase in C-to-U conversion rates compared to the un-crosslinked input sample (Figure 2D). We observed an additional 50 and 42 CIMS with significant cytosine to adenosine conversion in the day 7 and day 10 samples, respectively, bringing the total number of unique sites identified using m^6^A-eCLIP2 across both time points to 524. Notably, an adenosine is located at the -1 position of all identified cytosine sites, representing the putative methylated sites and confirming the specificity of this approach (Figure 2E). In addition, we find that the identified m^6^A CIMS are not equally distributed within transcripts but show a strong bias towards the 3’ UTR (Figure 2E), which is similar to the distribution pattern of m^6^A CIMS in other eukaryotes investigated^32,47^. Indeed, direct comparison of the read coverage of input and immunoprecipitated RNA across the transcriptome shows a higher ratio of immunoprecipitated RNA at the 3’ UTR, providing additional evidence for the unequal distribution of m^6^A CIMS within the gametocyte transcriptome. We therefore used an additional peak-calling approach to identify regions of putative m^6^A enrichment in the m^6^A-eCLIP2 data, identifying a total of 678 and 1,186 regions at day 7 and day 10, respectively.

A disadvantage of the m^6^A-eCLIP2 approach is its dependence on an m^6^A antibody that can have off-target affinities^32^, the indirect identification of m^6^A based on conversion rates downstream of the actual site, and its bias towards more abundant mRNAs. We therefore adapted GLORI-seq^48^, a chemical alteration assay in which un-methylated adenosines are converted to guanosines, which allows direct identification of m^6^A sites at single nucleotide resolution and calculation of relative methylation levels of each individual site^48^. This approach identified a total of 972 and 2,194 sites in day 7 and day 10 gametocytes, for a total of 2,349 unique sites in 1,159 transcripts across the gametocyte transcriptome.

The overall distribution of m^6^A localization identified using this approach mirrors that seen for m^6^A-eCLIP2, with 79% of sites (1,876/2,349 sites) located within the 3’ UTR or downstream of the coding sequence (Figure 2F,G). Interestingly, average methylation levels (i.e. the fraction of all transcripts transcribed from a given locus that is m^6^A methylated at a given site) reaches 64%. This is higher than that measured in mammalian cells^48^ and does not significantly differ between the two sampled timepoints during gametocytogenesis (Figure 2H).

Despite not relying on a downstream cytosine conversion as in m^6^A-eCLIP2, the GLORI-seq approach also identified a strong enrichment for cytosine at the +1 position of the m^6^A site (Figure 2F) and an overall similar RAC (R=A/G) motif that is observed in humans^29^ or yeast^32^. The preference for an adenosine at the -1 position in *P. falciparum* mirrors the substantially higher AU content found in the parasite’s genome, especially in UTRs (< 90%). Of note, we also find that methylation levels substantially differ depending on the motif, with non-RAC motifs showing a significantly lower methylation levels than RAC motifs (Figure 2I). Integration of the data from eCLIP2 and GLORI-seq (see Materials and Methods) resulted in 1,203 and 2,408 high confidence m^6^A sites identified at day 7 and day 10, respectively, totaling 2,743 unique m^6^A sites located in 1,296 transcripts across the *P. falciparum* gametocyte transcriptome (Figure 2J).

To determine if m^6^A is more predominant in male or female gametocytes, we categorized m^6^A-methylated transcripts into female- or male-dominant based on the gametocyte sex in which they were more highly expressed (see Materials and Methods). In total, we identified 589 and 1,300 m^6^A sites located in male- and female-dominant transcripts, respectively. Given that female cells account for ∼80% of all mature gametocytes, this difference can be attributed to overall higher sequencing coverage of female-dominant transcripts and therefore more stringent discovery in bulk transcriptome samples.

However, we identified similar numbers of m^6^A sites per transcript between the sexes (male: 1.9; female: 2.3), and the average methylation level does not differ significantly between male- and female-dominant, m^6^A-methylated transcripts (Figure 2K). Altogether, these data suggest a widespread m^6^A methylation program during gametocytogenesis that affects a range of biological process, but is largely agnostic to gametocyte sex.

### YTH.2 predominantly binds mRNA in male gametocytes

Although m^6^A methylation itself is present in both *P. falciparum* sexes, the YTH.2 reader protein is predominantly expressed in male gametocytes, and knockdown specifically affects male gametocyte activation. Thus, we next sought to characterize the interaction between m^6^A and YTH.2 in a sex-specific manner across gametocytogenesis. Males account for <30% of all gametocytes present in human blood cultures. In addition, protein and RNA input yields from cultured gametocytes are generally low, making the use of RNA-protein co-immunoprecipitation approaches for the identification of YTH.2. binding sites extremely difficult. We therefore developed an inducible protein-RNA proximity labelling approach based on surveying targets by APOBEC-mediated profiling (STAMP)^49^. Here, a cytosine-to-uridine (C-to-U) RNA base editor (i.e. APOBEC) was fused to the C-terminus of YTH.2, which allows for the editing of RNA nucleotides in close proximity to YTH.2 binding sites on mRNA transcripts (Figure 3A). Following standard RNA sequencing, regions of m^6^A-YTH.2 interactions can then be identified even from low input amounts based on signature C-to-U conversions.

**Figure 3:**
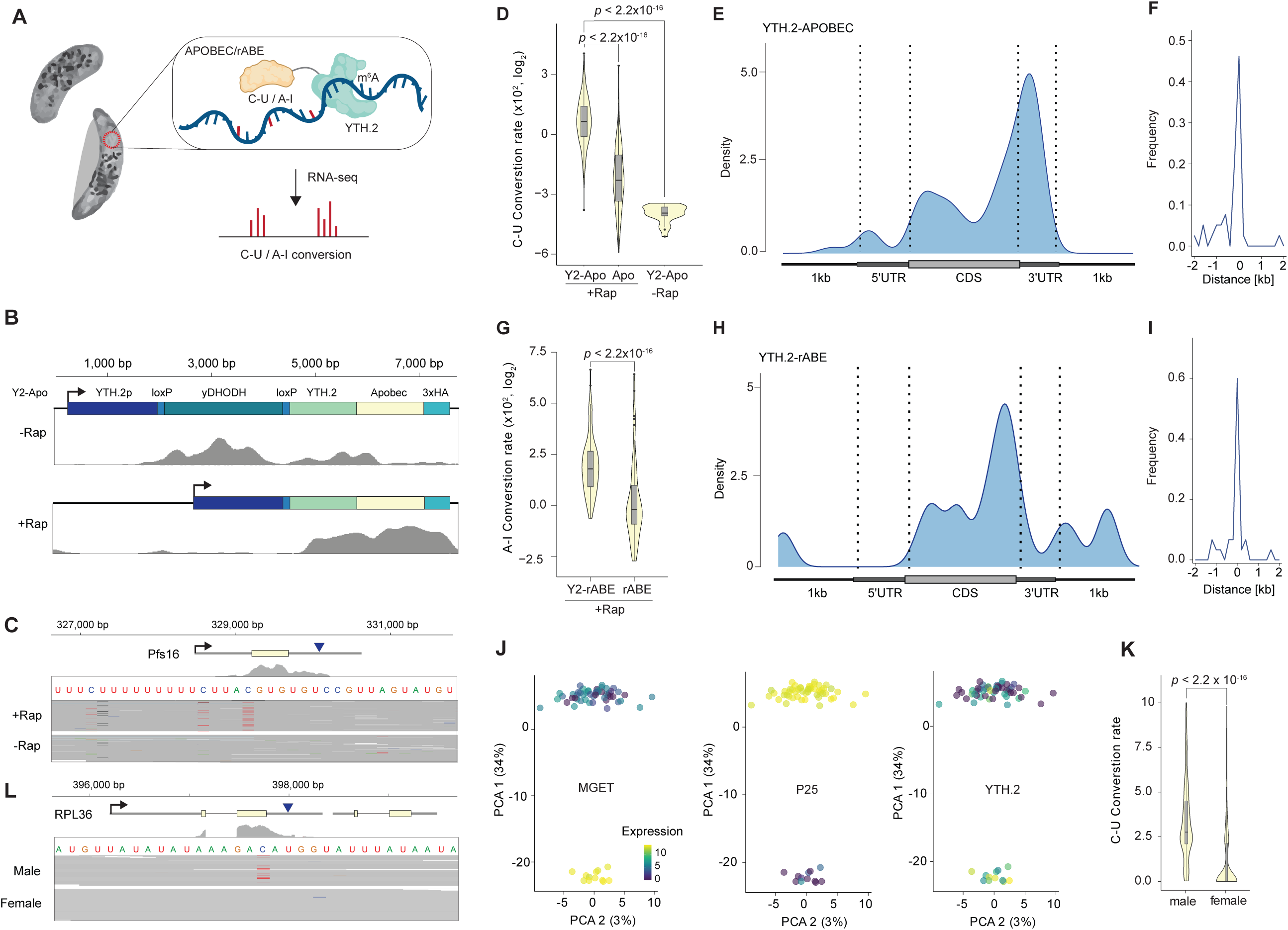
YTH.2 predominantly binds mRNA in male gametocytes. (A) Schematic of the YTH.2 protein-RNA proximity labelling approach. Sites with C-to-U (for APOBEC) or A-to-I (for rABE) conversions indicate regions of YTH.2 binding. (B) Expression construct and RNA-seq coverage for the inducible expression of YTH.2-APOBEC before treatment with rapamycin (-Rap, top) and after (+Rap, bottom). The RNA-seq coverage indicates precise excision of the yDHODH locus and activation of YTH.2-APOBEC transcription following rapamycin treatment (bottom). (C) Example of a C-to-U conversion detected in the induced (top) but not the un-induced YTH.2-APOBEC sample located in Pfs16. C-to-U conversions are indicated in red within individual aligned reads (grey). (D) Average conversion rates of high-confidence C-to-U sites identified in induced YTH.2-APOBEC cells (left) and corresponding conversion rates in APOBEC-only (center) and un-induced YTH.2-APOBEC. *p-* value were calculated using an unpaired Wilcoxon Rank Sum test. (E) Metagene plot representing the global distribution of high-confidence C-to-U conversions within transcripts detected by YTH.2-APOBEC. (F) Frequency plot showing the distance of high-confidence C-to-U conversion sites to the nearest m^6^A site. (G) Average conversion rates of high-confidence I-to-A sites identified in induced YTH.2-rABE cells (left) and corresponding conversion rates in rABE-only (right). *p-*value were calculated using an unpaired Wilcoxon Rank Sum test. (H) Metagene plot representing the global distribution of high-confidence A-to-I conversions within transcripts detected by YTH.2-rABE. (I) Frequency plot showing the distance of high-confidence I-to-A conversion sites identified by YTH.2-rABE to the nearest C-to-U conversion site identified by YTH.2-APOBEC. (J) Expression levels of MGET (left) and P25 (center) determined by ssRNA-seq of induced YTH.2-APOBEC cells, showing two distinct mature, male and female gametocyte populations. Expression of YTH.2 is shown on the right. (K) Conversion rates of high-confidence C-to-U sites identified in male (left) and female cells (right). *p-*value were calculated using an unpaired Wilcoxon Rank Sum test. (L) Example of a C-to-U conversion detected in male (top) but not female (bottom) YTH.2-APOBEC cells. C-to-U conversions are indicated in red within individual aligned reads (grey).

To inducibly express YTH.2-APOBEC, we generated an episome on which a *loxP*-flanked drug selection marker in inserted between the native *yth.2* promoter and a YTH.2-APOBEC coding region (Figure 3B, top). Transfection of this episome into a parasite cell line expressing dimerizable Cre recombinase allowed for the precise excision of the resistance maker after Cre activation by rapamycin and subsequent fusion of the upstream *yth.2* promoter to *yth.2-apobec* (Figure 3B, bottom). We activated Cre recombinase specifically at the onset of gametocytogenesis, leading to YTH.2-APOBEC expression. The same cell line without activation of Cre was used as a first negative control. In addition, we also transfected the same parent line with an episome with inducible expression of only APOBEC as a second negative control. Assaying C-to-U conversion led to the reproducible identification of 202 sites across the gametocyte transcriptome, with significantly higher C-to-U conversion than in either negative control (Figure 3C,D). Importantly, the identified YTH.2 binding regions show a predominant location within the 3’UTR, similar to m^6^A sites themselves (Figure 3E), and are centered around identified m^6^A sites (Figure 3F).

To further confirm these findings, we next used the same expression construct but replaced APOBEC with rABE, an adenine-to-inosine (A-to-I) base editor^50^. This approach identified a total of 138 reproducible A-to-I editing sites (Figure 3G) that show a similar 3’UTR-biased localization to m^6^A (Figure 3H). In addition, A-to-I editing sites are centered around the previously identified C-to-U editing sites using YTH.2-APOBEC1 (Figure 3I), confirming the reproducibility of the two independent approaches and identifying the *in vivo* binding of YTH.2 to cognate transcripts.

We next attempted to resolve a possible sex-specific binding of YTH.2 using YTH.2-APOBEC proximity labelling on a single cell level. To do so, YTH.2-APOBEC expression was activated at the onset of gametocytogenesis, and mature gametocytes were sorted based on sex using male- and female-specific markers and fluorescence-activated cell sorting. Full-length transcript single-cell RNA-seq was performed using FLASH-seq^51^, confirming two clear populations of male and female gametocytes (Figure 3J). To more stringently identify C-to-U editing sites between sexes, single-cell transcriptomes were pooled into one male and female sample, increasing the overall coverage across transcripts. This approach identified a total of 566 C-to-U conversion sites in either male or female gametocytes. Strikingly, 425 of those sites (75%) were either found exclusively in males (n = 304) or had a higher conversion rate in males than in females (n = 121, Figure 3K,L). Thus, the more pronounced binding of YTH.2 mirrors its predominant expression in male gametocytes, providing additional evidence for a sex-specific m^6^A methylation program.

### Male and female gametes engage in opposing translational programs during transmission

Given the RNA-binding properties of YTH.2^35,39^, we next investigated the effect of YTH.2 disruption on the gametocyte transcriptome. To avoid confounding the direct effects of YTH.2 depletion with broader effects related to parasite death following transmission, we compared NF54-YTH.2^WT^ to NF54-YTH.2^KD^ parasites. Only 22 genes were significantly altered in mRNA abundance in mature gametocytes, mirroring the limited general differences in expression pattern observed for select male and female marker genes (Figure 1E). Indeed, there were no clear differences in either male- or female-specific transcripts. Thus, YTH.2 does not appear to play a role in regulating mRNA transcript abundance in gametocytes.

As YTH.2 can repress mRNA translation during the asexual replicative cycle^39^, we next determined its effect on translation efficiency in gametocytes. To define a baseline of translational changes occurring during parasite transmission, we first characterized the translational trajectories of mRNA in bulk NF54-YTH.2^WT^ gametocytes. Using ribosome profiling as a direct approach for determining active protein synthesis in gametocytes, we found that female-dominant transcripts show overall lower translation efficiencies, confirming earlier findings of expansive translational repression during female gametocyte development on a transcriptome-wide level^19,21,28^. In contrast, we find that, on average, male-dominant transcripts have significant two-fold higher rates of translational efficiency than female-dominant transcripts (Figure 4A), suggesting that translational repression is not a feature of the male semi-quiescent transmission stage.

**Figure 4:**
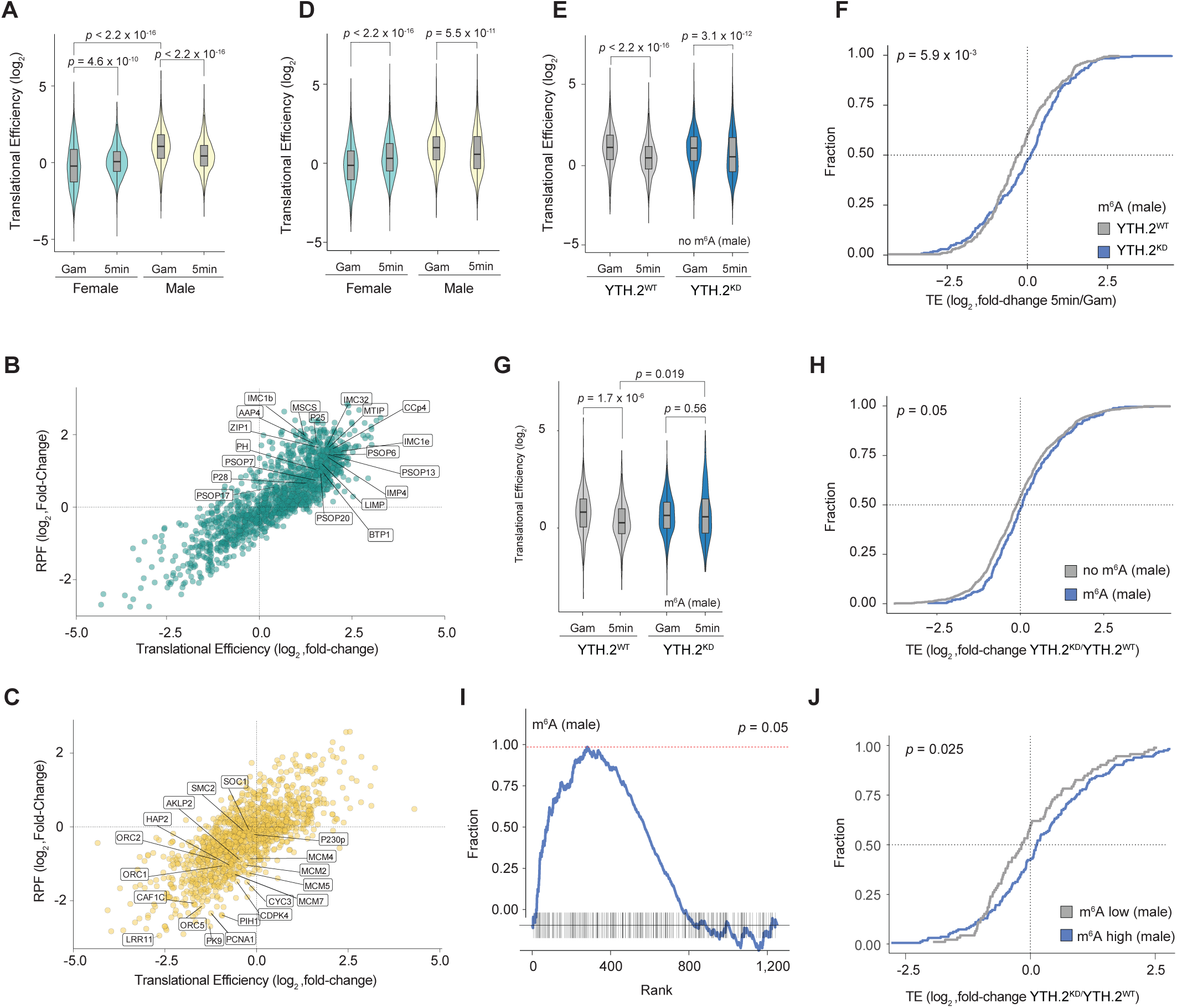
YTH.2 is essential for the translational shutdown of male-dominant m^6^A-methylated transcripts. (A) : Comparison of translational efficiencies of female- and male-dominant transcripts in mature NF54-YTH.2^WT^ gametocytes (Gam) and five minutes after transmission (5min). *p-*values were calculated using an unpaired Wilcoxon Rank Sum test. (B) Log_2_ fold-changes of ribosome footprints (RPF, ordinate) and translational efficiency (TE, abscissa) for female-dominant transcripts between mature gametocytes and 5min after transmission. (C) As I (B), but for male-dominant transcripts. (D) Translational efficiencies of female- and male-dominant transcripts in mature NF54-YTH.2^KD^ gametocytes (Gam) and five minutes after transmission (5min). *p-*values were calculated using an unpaired Wilcoxon Rank Sum test. (E) Translational efficiencies of non-m^6^A-methylated, male-dominant transcripts in mature NF54-YTH.2^WT^ (left) and NF54-YTH.2^KD^ (right) gametocytes (Gam) and five minutes after transmission (5min). *p-*value were calculated using an unpaired Wilcoxon Rank Sum test. (F) Cumulative fraction plot showing the proportion of m^6^A-methylated, male-dominant transcripts that feature increased or decreased translational efficiencies between mature gametocytes and 5min after transmission in NF54-YTH.2^WT^ (grey) and NF54-YTH.2^KD^ (blue) parasites. Following YTH.2 depletion. *p-* value was calculated using a two-sample Kolmogorov-Smirnov test. TE: Translational Efficiency (G) Translational efficiencies of m^6^A-methylated, male-dominant transcripts in mature NF54-YTH.2^WT^ (left) and NF54-YTH.2^KD^ (right) gametocytes (Gam) and five minutes after transmission (5min). *p-*value were calculated using an unpaired Wilcoxon Rank Sum test. (H) Cumulative fraction plot showing the proportion of male-dominant, m^6^A-methylated (blue) and non-m^6^A-methylated (grey) transcripts with increased or decreased translational efficiencies between NF54-YTH.2^WT^ and NF54-YTH.2^KD^ parasites five minutes after transmission. m^6^A methylated transcripts are more often upregulated five minutes after transmission when YTH.2 is depleted. *p-*value was calculated using a two-sample Kolmogorov-Smirnov test. TE: Translational Efficiency (I) Gene set enrichment analysis showing a significant enrichment of m^6^A methylation in male-dominant transcripts that are upregulated in NF54-YTH.2^KD^ compared to NF54-YTH.2^WT^ parasites five minutes after transmission. (J) Cumulative fraction plot showing the proportion of male-dominant, m^6^A-methylated transcripts with increased or decreased translational efficiencies in NF54-YTH.2^KD^ parasites five minutes after transmission depending on their methylation level. m^6^A methylated transcripts with higher m^6^A methylation levels (> 0.5) are more often upregulated than those with low m^6^A methylation levels (< 0.5) minutes after transmission when YTH.2 is depleted. *p-*value was calculated using a two-sample Kolmogorov-Smirnov test. TE: Translational Efficiency

We next performed ribosome profiling five minutes after the *in vitro* simulation of parasite transmission (i.e. after re-initiation of gametocyte development and during the formation of male and female gametes, see Materials and Methods). Comparing both the change in ribosome occupancy and translational efficiency (see Materials and Methods) showed that within this short timeframe, female-dominant transcripts immediately reverse translational repression and engage in extensive protein synthesis (Figure 4B), leading to a significant increase in overall translational efficiency (Figure 4A). Many female-dominant transcripts previously suggested to be translationally repressed prior to transmission, such as those encoding secreted membrane proteins, show a significant increase in ribosome occupancy upon ‘transmission’ (Figure 4B). In stark contrast, male-specific transcripts show the inverse pattern of translation, undergoing a significant decrease in both ribosome occupancy and translational efficiency upon transmission (Figure 4A,C). Such male-dominant transcripts include those encoding for proteins essential for DNA replication, nuclear division, and gamete fusion (Figure 4C). These data suggest that male gametocytes prepare for the rapid developmental progression following transmission by continuously transcribing and translating the proteins needed immediately after mosquito uptake. Once gamete differentiation is initiated, these transcripts are targeted for translational repression.

### YTH.2 is essential for the translational shutdown of male-dominant, m^6^A-methylated transcripts

To determine if YTH.2 plays a role in the translational transitions that takes place after gametocyte transmission, we next performed ribosome profiling on mature gametocytes and five minutes after *in vitro* transmission in NF54-YTH.2^KD^ parasites. In mature gametocytes, YTH.2 knock-down had no significant effect on overall translational efficiencies in male or female-dominant transcripts. Furthermore, upon transmission, YTH.2 knockdown did not affect the general patterns in translational efficiencies of female- or male-dominant transcripts (i.e. an increase in translational efficiencies in female- and a decrease in male-dominant transcripts) (Figure 4D). Thus, YTH.2 is not a general regulator of transcript translation.

Since YTH.2 is expressed predominantly in male gametocytes and specifically binds to m^6^A, we therefore explored sex-specific changes in translation efficiency of m^6^A-methylated transcripts. Indeed, when classifying transcripts based on their absolute levels of m^6^A methylation (as measured by GLORI-seq) in NF54-YTH.2^WT^ parasites, we found that the translational efficiency is more often downregulated for transcripts with higher m^6^A levels than for those with lower m^6^A levels during transmission, a trend that is especially true for male-dominant transcripts. In addition, we found no effect on the release from translational repression of female-dominant, m^6^A-methylated or non-methylated transcripts. Similarly, YTH.2 depletion has no effect on the decrease in translational efficiency of male-dominant, non-methylated transcripts (Figure 4E). However, male-dominant, m^6^A-methylated transcripts show a significant decrease in the number of transcripts whose translational is shut down five minutes after transmission in NF54-YTH.2^KD^ parasites. (Figure 4F). Concomitantly, overall translational efficiencies are maintained (rather than decreased) upon YTH.2 depletion five minutes after transmission at the same level as in gametocytes (Figure 4G), confirming that YTH.2 specifically effects translational repression of male-dominant, m^6^A-methylated transcripts.

We next directly compared translational efficiencies between NF54-YTH.2^WT^ and NF54-YTH.2^KD^ five minutes after transmission. Here, the translational efficiencies of m^6^A-methylated, male-dominant transcripts are significantly more often upregulated than those of non-methylated, male-dominant transcripts (Figure 4H), leading to a significant increase in average translational efficiencies of male-dominant, m^6^A-methylated transcripts (Figure 4E). Concomitantly, gene set enrichment analysis showed that male-dominant, m^6^A-methylated transcripts are significantly enriched among transcripts with an upregulation of translational efficiencies (Figure 4I), which is not seen for m^6^A-methylated, female dominant transcripts. Importantly, we find that this upregulation is further dependent on the absolute m^6^A methylation level (as measured by GLORI-seq) and specific to male gametocytes, where translational efficiencies of transcripts with higher m^6^A levels are significantly more often upregulated in NF54-YTH.2^KD^ parasites than those with lower m^6^A methylation levels five minutes after transmission (Figure 4J). As such, the efficient translational shutdown of m^6^A-methylated transcripts is specific to male gametocytes and dependent upon YTH.2.

### YTH.2 interaction with m^6^A-methylated transcripts is temperature-dependent

YTH.2 and m^6^A are readily detectable in mature gametocytes, yet translational efficiencies only decrease in a YTH.2-dependent manner after transmission, suggesting that an additional mediator or signal is necessary for the regulatory effect of YTH.2 on m^6^A-methylated transcripts. Extensive phosphorylation signaling cascades occur within the first minute following transmission in rodent malaria parasites^14,17,18^. However, the YTH.2 ortholog shows no change in phosphorylation during transmission, suggesting that this post-translational modification does not mediate m^6^A-YTH.2 interactions. Alternatively, temperature changes can substantially alter macromolecular interactions^52–58^, opening a possibly direct route for the restructuring of protein-RNA interactions. We therefore sought to decipher whether the temperature drop occurring during parasite transmission itself could alter m^6^A-YTH.2 interactions.

Measuring association and dissociation kinetics using surface plasmon resonance showed that interactions between YTH.2 protein^39^ and RNA is directly linked to the number of m^6^A sites present in the cognate RNA, with more m^6^A sites per RNA molecule leading to overall higher numbers and more stable YTH.2-RNA interactions (Figure 5A). We next tested the affinity of YTH.2 for RNA with a single m^6^A site at three different temperatures in a range that the parasite might experience during transmission. Strikingly, we find that the amount of YTH.2 interacting with m^6^A-methylated RNA at a given RNA concentration is substantially lower at 37°C (temperature of the human host) than at 32°C or 26°C (Figure 5B). Accordingly, the affinity of YTH.2 to m^6^A significantly increases over 10-fold (i.e. a decrease in equilibrium constant *K_D_*) at lower temperatures (Figure 5C), suggesting that the temperature drop during host-to-vector transmission itself alters m^6^A-YTH.2 interactions and could contribute significantly to translational shutdown.

**Figure 5:**
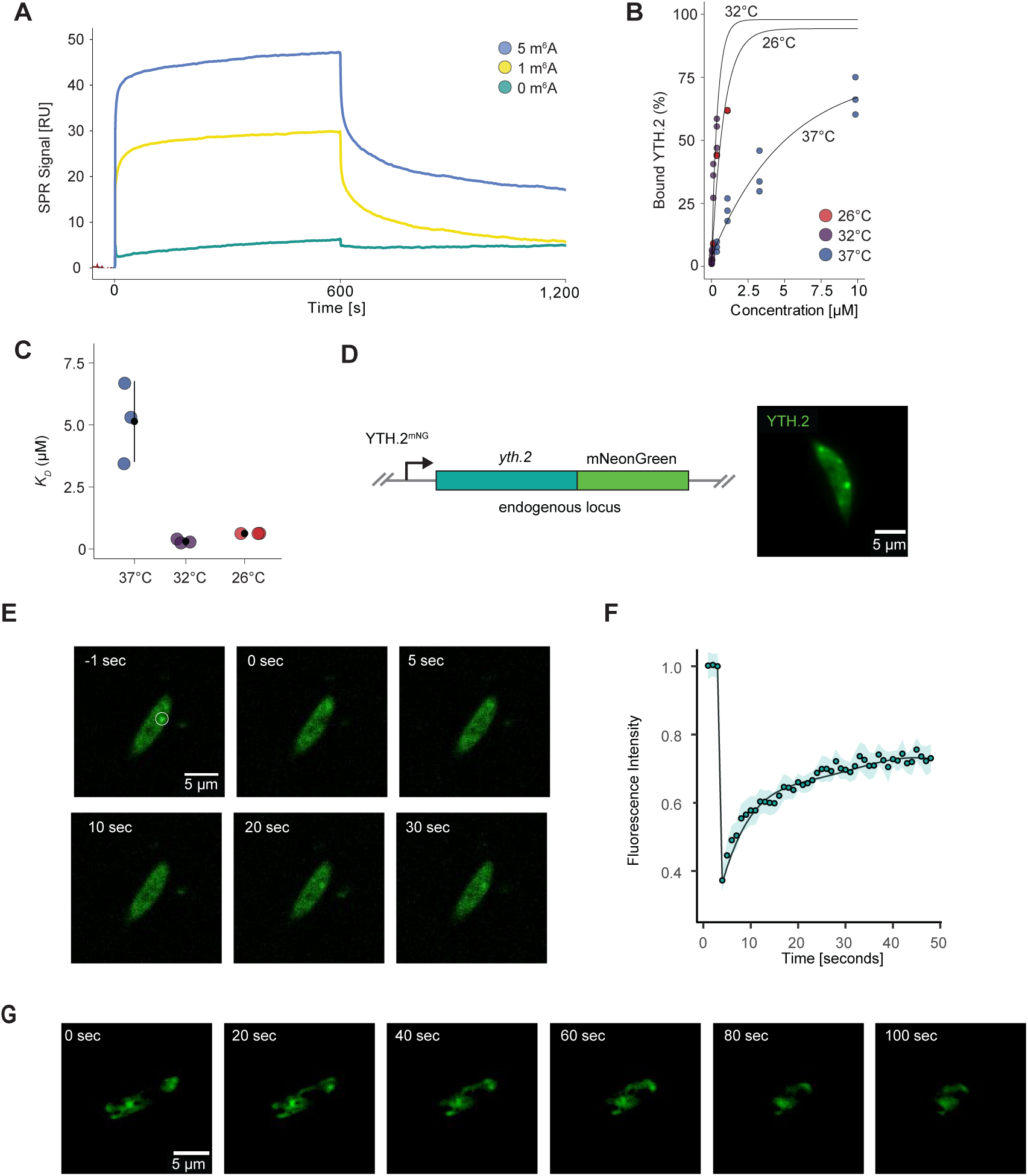
YTH.2 interaction with m^6^A transcripts is temperature-dependent. (A) Real time association and dissociation profiles corresponding to the injection of RNA with different numbers of m^6^A-methylated sites over immobilized GST-YTH.2 at 37°C. (B) Concentration dependence of YTH.2-RNA interactions (% of bound YTH.2, i.e. R_eq_/R_max_) at different temperatures. The synthetic RNA contains one m^6^A site. (C) Comparison of the equilibrium constant (*K_D_*) of YTH.2 interacting with RNA (one m^6^A site) at different temperatures. Black dot: mean; vertical line: standard error. Smaller *K_D_* indicates higher affinity. (D) Schematic of the native *yth.2* genic locus C-terminally fused with a mNeonGreen tag (mNG, left). Life-microscopy of YTH.2^mNG^ in mature gametocytes showing distinct cytoplasmic foci of higher YTH.2 concentration. (E) Example of fluorescence recovery of a cytoplasmic YTH.2^mNG^ foci after photobleaching. The bleached target region is indidcated with a white circle. (F) Quantification of the fluorescence recovery after photobleaching of cytoplasmic YTH.2^mNG^ foci. (G) Dynamics of YTH.2^mNG^ foci during the re-initiation of development after *in vitro* gametocyte activation.

In many eukaryotic cell types, translational repression coincides with the sequestration of mRNA into specific sub-cellular compartments (e.g. stress granules or P-bodies), making them inaccessible to translating ribosomes. To investigate how the m^6^A-mediated repressive effect on translation comes into effect following male gametocyte transmission, we followed YTH.2 protein localization in live gametocytes. We first fused *yth.2* with a sequence encoding mNeonGreen at the endogenous genomic locus (Figure 5D). Live imaging of mature gametocytes showed that YTH.2 forms specific foci within the cytoplasm (Figure 5D). Using fluorescent recovery after photobleaching, we found that YTH.2 rapidly diffuses back into these foci, reaching 70% fluorescence recovery less than 50 seconds after photobleaching (Figure 5E,F). These data suggest that YTH.2 is highly mobile in mature gametocytes despite localizing to these membrane-less compartments. In contrast, within the first minutes after *in vitro* activation of gametocytes, we found that YTH.2 foci remain stable (Figure 5G). Hence, following increased m^6^A-YTH.2 affinities after transmission, YTH.2 foci might sequester m^6^A-methylated transcripts away from the translatable pool, suppressing protein synthesis.

## DISCUSSION

The rapid re-initiation of development and differentiation into gametes after transmission from the human host to its mosquito vector is a hallmark of human malaria parasites. However, the mechanisms by which male parasites prepare for the exit from their semi-quiescent gametocyte stage and subsequently rewire their transcriptome in response to the environmental changes accompanying transmission are poorly understood. The data from this study suggest that the mRNA modification N6-methyladeosine and its reader protein YTH.2 prime the male gametocyte transcriptome for a rapid shutdown of protein synthesis and that their repressive interaction is mediated by the temperature decrease accompanying transmission.

We integrated two orthogonal, nucleotide-resolution m^6^A mapping approaches to directly identify and quantify m^6^A methylation transcriptome-wide in mature *P. falciparum* gametocytes. Interestingly, we find a strong 3’-biased localization of m^6^A in *P. falciparum* transcripts as well as a conserved RAC motif that resemble patterns seen in model organisms. Importantly, this 3’ end bias in model systems has been attributed to the fact that m^6^A methylation is the default state of DRACH motifs and is excluded from most internal exons due to the collision of the methyltransferase complex and the spliceosome. However, most protein-coding genes (2,336/5,285), including those with an m^6^A site (510/1,296) in *P. falciparum* are single-exon genes, which rules out the possibility of a large effect of splicing. In addition, RAC motifs are substantially more abundant within coding regions of *P. falciparum* (CDS = 22.3/kb of CDS vs 12.4/kb of 3’ UTR). Hence, while certain MT-A70-interacting proteins might mediate the specificity of m^6^A methylation, the question of how m^6^A sites are specifically targeted to the 3’ UTR remains open.

m^6^A methylation itself is present in both male and female gametocytes and might regulate two distinct functions; A potential ‘housekeeping’ role in both sexes may be mediating correct 3’ end processing via the conserved YTH.1 protein^36–38^, which is expressed equally in both sexes. On the other hand, our data also revealed a male gametocyte-specific role in translational repression following transmission specifically via YTH.2, which is the opposite of the translational pattern seen in female gametocytes. Female gametocytes transcribe but extensively repress translation of mRNAs it requires only after transmission to the mosquito vector^19,22^. While translational efficiency is low in the human host, it is rapidly upregulated after mosquito uptake on a transcriptome-wide level. Stockpiling these mRNAs in specific subcellular compartments prior to mosquito uptake could allow the female gametocyte to ‘skip’ the time-consuming process of transcriptional activation and go straight to protein synthesis to achieve more rapid differentiation into a macrogamete.

Here, we show that male gametocytes use a similarly extensive post-transcriptional mechanism to prepare for transmission. Instead of repressing the translation of gamete-specific proteins, male gametocytes sustain high levels of translation for proteins it requires to immediately re-initiate development and engages in an extensive shutdown of translation once transmitted. Male gametocytes thus forego some transcriptional as well as post-transcriptional regulatory mechanisms of gamete-specific genes and stockpile key proteins, which may provide a higher level of ‘readiness’ during their semi-quiescent stage and preparation for transmission. The activation of proteins that regulate differentiation into male gametes might therefore be predominantly regulated on a post-translational level. Indeed, activation of male gametocytes depends on the activation of cyclic guanosine monophosphate (cGMP)– dependent protein kinase (PKG) that mobilizes Ca_2_^+^ and facilitates an extensive cascade of phosphorylation signaling within seconds following transmission^14,17,18^. Importantly, this short timeframe requires these proteins to already be available upon transmission. In fact, we find these proteins that are modified upon transmission to be among the most highly translated transcripts in male gametocytes.

’Skipping’ of some of the transcriptional and especially post-transcriptional regulation in gametocytes, however, requires an even more precise and rapid control of protein synthesis after transmission. Our data indicate that through m^6^A methylation of cognate transcripts and the expression of YTH.2, the male transcriptome is already prepared in gametocytes to rapidly shutdown translation upon transmission. Integrating a temperature decrease as the mediator of repressive m^6^A-YTH.2 interactions ensures that translational efficiencies only decrease after transmission. Still, m^6^A-YTH.2 interactions might represent only one of multiple mechanisms that facilitates this shutdown, given that translational efficiencies of non-methylated transcripts also decrease after transmission. Since YTH.2 is essential for male gamete differentiation, however, parallel repressive mechanisms likely act independently and cannot complement each other. Given that both interacting partners are already highly abundant, their interaction might allow for a much faster translational shutdown after the temperature drop before other mechanisms come into effect.

Sensing of environmental cues has already been reported to be essential for the activation of male gametocytes, especially by maintaining levels of the secondary messenger cGMP in gametocytes through temperature-sensitive phosphodiesterases and a pH- and XA-responsive membrane complex centered around the guanylyl cyclase GCα^14^. However, a temperature drop that promotes the interaction of macromolecular complexes and mediates a widespread post-transcriptional mechanism represents a more direct and rapid integration of an environmental signal. Our *in vivo* data on the dynamics of translational efficiencies suggests that a purely thermodynamic regulation leading to a non-specific, global increase in mRNA-protein affinities at lower temperatures is not solely sufficient to effect translational repression since female-dominant transcripts actually increase in translational efficiency during transmission. Moreover, YTH.2 alone is also not sufficient to repress protein synthesis, given that it is abundantly present in gametocytes and m^6^A-methylated transcripts are translated at higher rates in gametocytes than after transmission. Altogether, this indicates that only the combination of m^6^A, a temperature decrease, and YTH.2 can mediate the translational repression of m^6^A-methyalted transcripts following transmission. Interestingly, a temperature decrease of only 5°C is already sufficient to increase the affinity of YTH.2 to m^6^A-methyalted RNA by over an order of magnitude. However, affinity does not further increase at even lower temperatures, suggesting a temperature switch rather than a gradual temperature dependence of YTH.2-m^6^A interaction. The fact that mosquitoes preferentially feed at night might ensure that even if environmental temperatures (and thus the temperature of the mosquito) are close to 37°C, male parasites can still efficiently transmit.

Importantly, in addition to the presence of all three factors (i.e. m^6^A, YTH.2 and temperature), mediation of translational repression of m^6^A-methylated transcripts possibly further relies on the specific subcellular localization of YTH.2. While YTH.2 (and possibly RNA) rapidly diffuses into and out of membrane-less cytoplasmic compartments at 37°, it stabilizes after a temperature drop and the re-initiation of development. Therefore, m^6^A-methylated transcripts might remain within the translatable pool of transcripts in gametocytes and transiently interact with YTH.2 foci, but they are sequestered to these foci upon the temperature drop accompanying transmission and the simultaneous increase of m^6^A-YTH.2 affinities.

In conclusion, we identify a uniquely extensive and sex-specific epitranscriptomic mechanism that allows malaria parasites to prepare for transmission to the mosquito vector and concurrently ‘primes’ the transcriptome for a rapid rewiring of translational outputs. The integration of a temperature signal exemplifies how malaria parasites not only adapted to, but actively utilize sudden environmental changes to control molecular mechanisms during a major developmental transition.

## MATERIALS AND METHODS

### Parasite culture

Asexual blood-stage *P. falciparum* parasites were cultured as described previously^60^. In brief, parasites were cultured in human RBCs (obtained from the Etablissement Francais du Sang with approval number HS 2021-24819) in RPMI-1640 medium (Thermo Fisher # 53400-025) supplemented with 10% v/v Albumax I (Thermo Fisher no. 11020039), hypoxanthine (0.1 mM final concentration, CC-Pro # Z-41-M) and 10 mg gentamicin (Sigma # G1397-10ML) at 4% hematocrit and under 5% O2, 3% CO_2_ at 37°C. Development of parasites was monitored thin blood smears and Giemsa staining. To tightly synchronize the developmental stages of asexually replicating parasites, late stages were enriched by plasmion floatation followed by ring-stage enrichment using 5% sorbitol treatment 6 hours later. The 0 h timepoint was considered to be 3 hours after the plasmion treatment.

### Gametocyte induction

Synchronized gametocyte development was induced following the protocol of Fivelman et al.^61^ Synchronous asexual, late-stage parasites (∼35 h.p.i.) were concentrated at ∼2.5% parasitemia and 2.5% hematocrit using plasmion flotation. The next day, 75% of the spent culture medium was replaced with fresh medium, and the ring-stage parasites (∼10-15% parasitemia) were left to develop into trophozoites at high parasitemia for an additional 24h. The culture was then diluted in fresh media to 3% parasitemia and kept for an additional 24 hours. The growth medium of the resulting high-parasitemia, ring-stage culture (Day 0 of gametocyte development) was then replaced with RPMI supplemented with 5% human serum, 5% Albumax, 0.1 mM hypoxanthin, 10 mg gentamicin and 50 mM N-acetylglucosamine (NAG, Sigma # A3286). Growth media was then changed daily for 5 days with the addition of NAG to prevent growth of asexually replicating parasites, and then without NAG for an additional five days. Mature, stage V gametocytes were harvested at day 10.

### Total RNA extraction and mRNA enrichment

For all RNA-sequencing based methods, gametocytes of wild-type *P. falciparum* (strain NF54) were induced as described above. Red blood cells were lysed with 0.075% saponin in DPBS at 37°C, and the resulting parasite cell pellet was washed once with ice-cold DPBS and then resuspended in 700 μl QIAzol reagent (Qiagen # 79306). Total RNA was extracted using the Qiagen miRNeasy kit (Qiagen # 217004) including an on-column DNase I digestion according to the manufacturer’s protocol. mRNA was enriched using the Dynabeads mRNA purification kit (Thermo Fisher # 61006).

### Analysis of mRNA modifications by LC-MS/MS

Total RNA samples were collected, and mRNA was enriched as described above. The mRNA samples were then processed and analyzed as described previously^33^

### m^6^A-eCLIP2

Total RNA extraction and mRNA enrichment was performed as described above. For m^6^A-eCLIP2, a total of two samples were collected at day 7 and day 10 post gametocyte induction. RNA barcoding and pooling followed the protocol described by Dierks et al.^46^. In brief, for each individual sample the mRNA was fragmented using the Zinc RNA fragmentation kit (Thermo Fisher # AM8740) for 2.5 min at 70°C, resulting in an average length of 200nt and the reaction was cleaned up using RLT Buffer (Qiagen #79216) and Dynabeads MyOne Silane beads (Thermo Fisher # 370-02D). Next, contaminating DNA was removed, RNA fragments dephosphorylated and then 3’ phosphorylated in a single reaction by incubating the sample with TurboDNase (Thermo Fisher # AM2238), FastAP (Fermentas # EF0651) and T4 PNK (NEB # M0201L) for 30 mins at 37°C. Barcoded 3’ adapters were ligated to the RNA fragments using T4 RNA Ligase I and incubation for 1.5h at 23°C. The barcoded RNA was then cleaned up and pooled at equal ratios into a single sample. A sample serving as input control was put aside for later processing. Next, RNA was diluted in immunoprecipitation (IPP) buffer (150 mM NaCl, 0.1% NP40, 10 mM Tris-HCl) and incubated with anti-m^6^A antibody (Abcam # ab151230) for 2 h at 4°C. The antibody-coupled RNA was cross-linked twice at 254 nm with 150 mJ/cm^2^ in a UV-crosslinker (Thermo Fisher # 16556804), and then immunoprecipitated for 1h at 4°C using protein G magnetic beads (Thermo Fisher # 10003D). The beads were washed twice in IPP buffer, low salt buffer (50 mM NaCl, 0.1% NP40, 10 mM Tris-HCl), high salt buffer (500 mM NaCl, 0.1% NP40, 10 mM Tris-HCl) and then once in again in IPP buffer. The cross-linked RNA-antibody complexes were eluted from the protein G magnetic beads by mixing them with 1X Loading buffer (10uL 4X LDS Sample buffer (Thermo Fisher # NP0007) with 30uL + 1X IPP buffer) and incubation at 70°C for 10 minutes with interval mixing at 1,200 rpm. The supernatant was separated by size on a NuPAGE 4-12% Bis-Tris gel (Thermo Fisher # NP0321) and then transferred to a nitrocellulose membrane to remove unbound RNA. Membrane regions corresponding to the estimated size of cross-linked RNA-antibody complexes (30-150 kDa) were excised from the membrane. To recover the RNA from the membrane and remove the m^6^A antibody, the membrane was first incubated with Proteinase K for 20 min at 37°C and then at 50°C for an additional 20 min with interval mixing at 1,200 rpm. At this point, recovered RNA still retains covalently bound peptides from the antibody at the m^6^A site that can introduce signature substitutions during the next step^44^. RNA was reverse transcribed using Superscript III reverse transcriptase and primers were removed using ExoSap-IT (Affymetrix # 78201) following standard protocols. RNA was then hydrolyzed using NaOH and single-strand cDNA was cleaned up using RLT buffer and silane beads. 5’ adapters were ligated to the cDNA using T4 RNA Ligase 1 overnight at 23°C and then PCR-amplified using the KAPA HiFi HotStart ReadyMix with primers P5 Solexa/P3 Solexa. The final library was purified using AMPure XP beads (Beckman # A63880) and controlled for adapter-dimers and concentration on a Tapestation (Agilent) with a High sensitivity D1000 screen tape (Agilent # 067-5584).

### GLORI-seq

GLORI-seq was performed as described previously.^48^ In brief, mRNA was fragmented at 94°C for 3 min in NEB RNA 10X fragmentation buffer and cleaned up by ethanol precipitation. For RNA protection, the RNA was mixed with glyoxal, DMSO and RNase-free water and incubated for 30 min at 50°C. After 30 min incubation, saturated H_3_BO_3_ was added to the reaction and incubated for another 30 min at 50°C and placed on ice for 2 min after the second incubation time. For the deamination of the RNA, 5M NaNO_2_, 500mM MES (pH 6.0) and 8.8 M Glyoxal were added to the sample and incubated in a thermocycler for 8 hours at 16 °C. Subsequently, the RNA was cleaned up by precipitation, the pellet was resuspended in 50 μl deprotection buffer (10 ml 1M Triethylammonium pH 8.6, 9.5 ml deionized formamide, 0.5 ml RNase-free water) and incubated in a thermocycler at 95°C for 10 min. For end repair with T4 PNK, the reaction RNA from the previous step was mixed with 10X T4 PNK Buffer, T4 PNK, Murine RNase inhibitor and RNase-free water and incubated at 37°C for 30 min, then 10 mM ATP was added and the reaction was incubated for another 30 min. Afterwards, the RNA was cleaned once again using the Zymo Research RNA Clean & Concentrator Kit. RNA libraries where then generated using the NEBNext Small RNA Library Prep Kit for Illumina. The small RNA libraries were pooled and size-selected using a Pippin Prep (Sage Science). Final libraries were sequenced on a NextSeq2000 in a 2×151bp layout.

### Surface Plasmon Resonance

The experiments were carried out on a Biacore^TM^ T200 (Cytiva). For each experiment (different temperature), a new Series S Sensor Chip CM5 (Cytiva) and the reagents from the GST Capture kit by Cytiva were used.

Every experiment started by the immobilization of anti-GST antibody (10μg/ml) on a new Series S Sensor Chip CM5 (Cytiva) chip according to the manufacturer’s protocol. The buffer was changed from PBS (used for cleaning and immobilization) to the running buffer for the experiments (50mM Tris-HCl pH7.5, 150 mM NaCl). Unbound anti-GST was removed by flowing with GST (10μg/ml) for 60s at 5μl/min followed by 120s regeneration solution Glycine-HCl pH2.1. The first step of the actual SPR measurement was the capture of GST (10μg/ml, diluted in running buffer) in flow cell 1 (as a reference) resulting in 1128 RUs on average, followed by the capture of GST-YTH.2 (10μg/ml, diluted in running buffer) in flow cells 2, 3 and 4, one by one at 5μl/min, resulting in 2562 RUs on average. Afterwards, six injections (600s each, 10 μl/min) of running buffer with BSA (50mM Tris-HCl pH7.5, 150 mM NaCl, 1mg/ml BSA) were flowed to attain a stable base line signal before the injection of the different RNAs. The RNAs were diluted in the running buffer complemented with BSA (50mM Tris-HCl pH7.5, 150mM NaCl, 1mg/ml BSA) to prevent sticking to the sample tube. Flow paths 1-2-3-4 were opened to inject the RNAs for 600s at a flow rate of 10μl/min, followed by 600s of dissociation. In-between different RNA injections, the flow cells were regenerated with two injections of 4M NaCl to remove bound RNA from the protein before injecting a new concentration. The concentrations used in the experiments were chosen based on the lowest concentration giving a signal at the respective temperature. A random order of injections of the different RNAs and concentrations was chosen to prevent any bias. We injected five concentrations for each RNA with a 3-fold dilution factor between concentrations. At 26°C the concentration range for RNA with 0 m^6^A sites was 40.5nM to 3280.5nM, 1 m^6^A site 13.5nM to 1093.5nM and 5 m^6^A sites 0.5nM to 40.5nM. At 32°C the concentration range for RNA with 0 m^6^A sites was 13.5nM to 1093.5nM, 1 m^6^A site 4.5nM to 364.5nM and 5 m^6^A sites 0.17nM to 13.5nM. At 37°C the concentration range for RNA with 0 m^6^A sites was 1093.5nM to 88573.5nM, 1 m^6^A site 364.5nM to 29424.5nM and 5 m^6^A sites 13.5nM to 1093.5nM.

The interaction kinetics were determined from the change in response (=SPR signal, RU) as a function of time represented as sensograms (Figure 5A). The sensograms were reference subtracted and blank subtracted with the Biacore T200 Evaluation Software Version 3.2.1 and Version 2.0, and further analyzed with Scrubber Version 2.0 and BIAEVAL Version 4.0. As a fitting model for affinity, we used the steady state affinity model performed with the Biacore T200 Evaluation Software Version 3.2.1 and Version 2.0. The model calculates the equilibrium dissociation constant (K_D_) for a 1:1 interaction from a plot of steady state binding levels (R_eq_) against analyte concentration (C): 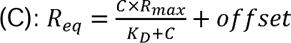 (Figure 5C). Percentage of YTH.2 bound was calculated as R_eq_/R_max_ (analyte binding capacity of the surface) and plotted over the RNA concentration with BIAEVAL Version 4.0 (Figure 5B).

